# Drivers and Determinants of Strain Dynamics Following Faecal Microbiota Transplantation

**DOI:** 10.1101/2021.09.30.462010

**Authors:** Thomas SB Schmidt, Simone S Li, Oleksandr M Maistrenko, Wasiu Akanni, Luis Pedro Coelho, Sibasish Dolai, Anthony Fullam, Anna M Glazek, Rajna Hercog, Hilde Herrema, Ferris Jung, Stefanie Kandels, Askarbek Orakov, Thea Van Rossum, Vladimir Benes, Thomas J Borody, Willem M de Vos, Cyriel Y Ponsioen, Max Nieuwdorp, Peer Bork

## Abstract

Faecal microbiota transplantation (FMT) is an efficacious therapeutic intervention, but its clinical mode of action and underlying microbiome dynamics remain poorly understood. Here, we analysed the metagenomes associated with 142 FMTs, in a time series-based meta-study across five disease indications. We quantified strain-level dynamics of 1,089 microbial species based on their pangenome, complemented with 47,548 newly constructed metagenome-assembled genomes. Using subsets of procedural-, host- and microbiome-based variables, LASSO-regularised regression models accurately predicted the colonisation and resilience of donor and recipient microbes, as well as turnover of individual species. Linking this to putative ecological mechanisms, we found these sets of variables to be informative of the underlying processes that shape the post-FMT gut microbiome. Recipient factors and complementarity of donor and recipient microbiomes, encompassing entire communities to individual strains, were the main determinants of individual strain population dynamics, and mostly independent of clinical outcomes. Recipient community state and the degree of residual strain depletion provided a neutral baseline for donor strain colonisation success, in addition to inhibitive priority effects between species and conspecific strains, as well as putatively adaptive processes. Our results suggest promising tunable parameters to enhance donor flora colonisation or recipient flora displacement in clinical practice, towards the development of more targeted and personalised therapies.

## Introduction

Faecal microbiota transplantation (FMT) involves the transfer of gut microbes, viruses and luminal content to modulate a recipient’s microbiome, usually for therapeutic purposes. While the efficacy of FMT has been demonstrated for various diseases ^1–3^, such as recurrent *Clostridioides difficile* (rCDI) infection ^4–6^, ulcerative colitis (UC ^7,8^) or cancer cachexia ^9^, it may also facilitate microbiome recovery following disturbance ^10^ and can enhance microbiome-mediated responses to other therapies ^11,12^. Yet despite demonstrable efficacy in a growing range of clinical applications, the mode of action of FMT remains poorly understood ^3,13^, and neither clinical success nor adverse outcomes are currently predictable with accuracy.

As FMT primarily targets the microbiome, the engraftment of ‘beneficial’ and/or displacement of ‘detrimental’ microbes are expected to cause clinical effects ^3^, in conjunction with more specific processes of host-microbiome interplay, such as the modulation of immune responses ^14^, restored short chain fatty acid (SCFA) metabolism ^13^ or reinstated phage pressure ^15,16^. It has been argued that both microbiome engraftment and clinical success are mainly determined by donor factors, and that rationally selected ‘super-donors’ may improve therapeutic efficacy ^17,18^. This donor-centric view has since been questioned at least for some indications ^19^, highlighting the importance of recipient ^20^ or procedural ^21^ factors instead.

Changes in microbial compositions following FMT have been studied with regard to phages ^22^ or fungi ^23,24^, yet the bulk of current knowledge is focussed on bacteria and archaea, where colonisation by donor microbes and the persistence of indigenous recipient microbes emerge at the strain level of microbial populations ^25^. Strain-level studies suggest that colonisation levels following FMT vary across indications: whereas donor and recipient strains coexist long-term in metabolic syndrome (MetS) patients ^25^, donor takeover is the most common outcome in rCDI ^26– 29^, with intermediate outcomes in UC ^30^ or obesity ^31^. However, the factors shaping these differential strain-level outcomes remain poorly understood. In small pilot study cohorts, colonisation success of donor strains leading to short-term persistence was associated with species phylogeny, broad microbial phenotypes and relative faecal abundances in rCDI ^26,27^, but with more adaptive metabolic phenotypes in UC ^32^.

Here, we conducted a meta-analysis of FMT strain-level outcomes across multiple indications, combining newly generated with public data. We hypothesised that drivers of FMT response are best studied from an ecological perspective ^33,34^: FMTs can be thought of as untargeted perturbation experiments on the gut microbiome *in natura*, pitting donor communities against recipient, whose outcomes emerge from underlying ecological processes. We therefore quantified strain-level patterns of donor colonisation, recipient resilience and turnover following FMT, both at the broad level of entire communities and specifically for individual species. We built cross-validated models to predict FMT outcome based on either *ex ante* variables (i.e. knowable before the intervention) or *post hoc* readouts (measured after the intervention), further categorised by scope (procedural, donor-related or recipient-related) and resolution (host-, community-, strain-level), yielding testable hypotheses. Linking informative variables and their predictive performance to putative underlying ecological processes, we provide a comprehensive view of host- and microbiome-level determinants of strain dynamics following FMT, with relevance to gut microbial ecology in the clinical context and beyond.

## Results

### A meta-analysis of FMT strain-level outcomes across indications

We analysed 556 faecal metagenomes collected in 142 time series of FMTs conducted for ulcerative colitis (UC, n=42 ^30,35–37^), recurrent *Clostridioides difficile* infection (rCDI, n=37 ^26,28,38^), metabolic syndrome (MetS, n=26 ^25,39^), infection with extended-spectrum beta lactamase producing bacteria (ESBL, n=19 ^40^), or Crohn’s disease (CD, n=18 ^41^). The majority (n=121) of analysed time series were allogenic FMTs (transfer of donor stool to a recipient), the remainder (n=21) were autologous transfers (of the recipient’s own stool); see tables S1-3 and Methods for a full dataset description. We profiled 1,089 prevalent microbial species, including 144 that were previously undescribed, in meta-pangenomes constructed from 47,548 newly built metagenome-assembled genomes (MAGs) and 25,037 reference genomes ^42^ (Fig 1A; Methods).

**Figure 1.**
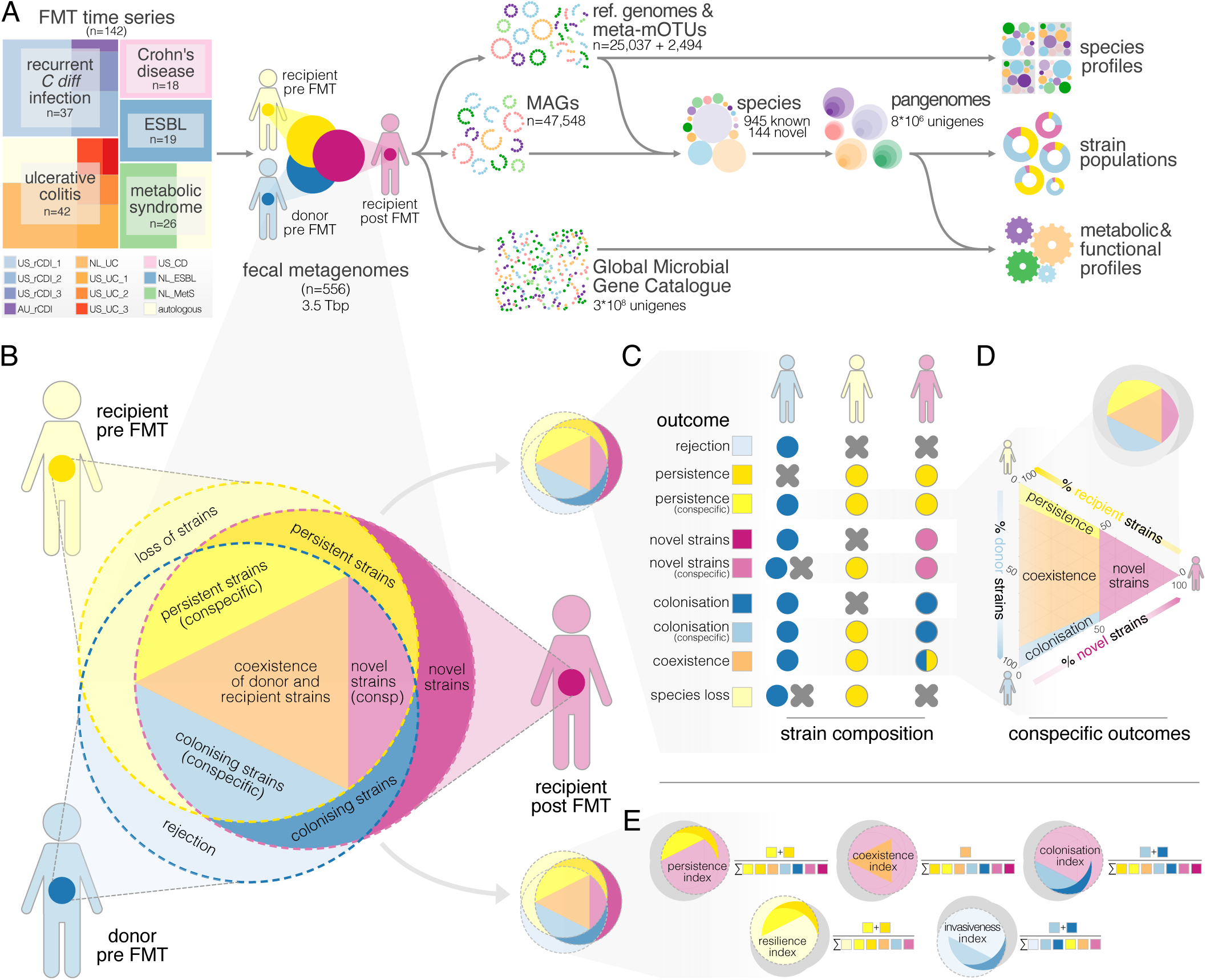
Study design & workflow overview. (A) We analysed a dataset of 142 FMT time series across five indications and 12 cohorts, totalling 556 faecal metagenomes. Species pangenomes were built from reference genomes and newly generated MAGs and profiled across samples for taxonomic, functional and strain population composition, based on microbial single nucleotide variants (SNVs) and differential gene content. (B) Each FMT was represented as a triplet of donor pre-FMT (blue hues), recipient pre-FMT (yellow) and post-FMT (purple) samples; each sample’s strain population is indicated as an overlapping circle. (C) FMT strain-level outcomes for each species were scored using patterns of determinant SNVs and gene content; see table S5 for details. (D) Ternary diagram of the strain population space for conspecific persistence, colonisation, coexistence and influx of novel strains. (E) Outcomes across all observed species in a sample triplet were further summarised using persistence, resilience, coexistence, colonisation and invasiveness indices (see Methods).

Based on observed differences in determinant single nucleotide variants (SNVs) and gene content in donor and recipient baseline microbiomes, and changes in recipient post-FMT samples (Fig 1B), we classified patterns in post-FMT strain population composition for each identified species as: recipient *persistence* (population dominated by recipient strains), donor *colonisation* (dominated by donor strains), donor-recipient strain *coexistence*, influx of novel or previously undetected strains, donor *rejection* (failure to colonise beyond detection limit) and *loss* of all recipient strains (Fig 1B-D; Methods, table S5). Based on these categories, we calculated summary statistics for FMT outcomes across species (Fig 1E; Methods): a *persistence, coexistence* and *colonisation index* as the total fraction of recipient-specific, coexisting and donor-specific strains in the post-FMT microbiome; a *resilience index* as the fraction of recipient strains persisting after FMT; and an *invasiveness index* as the fraction of donor strains that successfully engrafted.

### High inter-individual variability of strain-level outcomes following FMT, independent of clinical effects

Strain-level outcomes varied greatly among allogenic FMTs when summarised across all tracked species (Fig 2A). We did not observe complete recipient strain turnover (loss of all strains) or complete donor rejection (failure to colonise) in any analysed FMT, although recipient persistence or donor colonisation were very low in some patients. Outcome generally varied based on pre-FMT species occupancy: both donor takeover (accounting for 20.1±18.1% species post-FMT) and recipient persistence (11.2±9.2%) were more frequent among species with complementary pre-FMT occupancy, i.e. presence in either donor or recipient, but not both. In contrast, conspecific coexistence of donor and recipient strains (21.6±14.2%) was the most frequent outcome for species present in both donor and recipient pre FMT, compared to conspecific colonisation (3.7±3.1%) and persistence (5.5±5.6%). More than a third (37.8±11.5%) of post-FMT strain populations were attributable to novel strains or entirely novel species, i.e. not present in either donor or recipient pre-FMT, or previously below detection limit. Such major turnover towards novel strains was likely associated with the intervention itself, as novel strains accounted for 53±11% in autologous (‘placebo’) FMTs.

**Figure 2.**
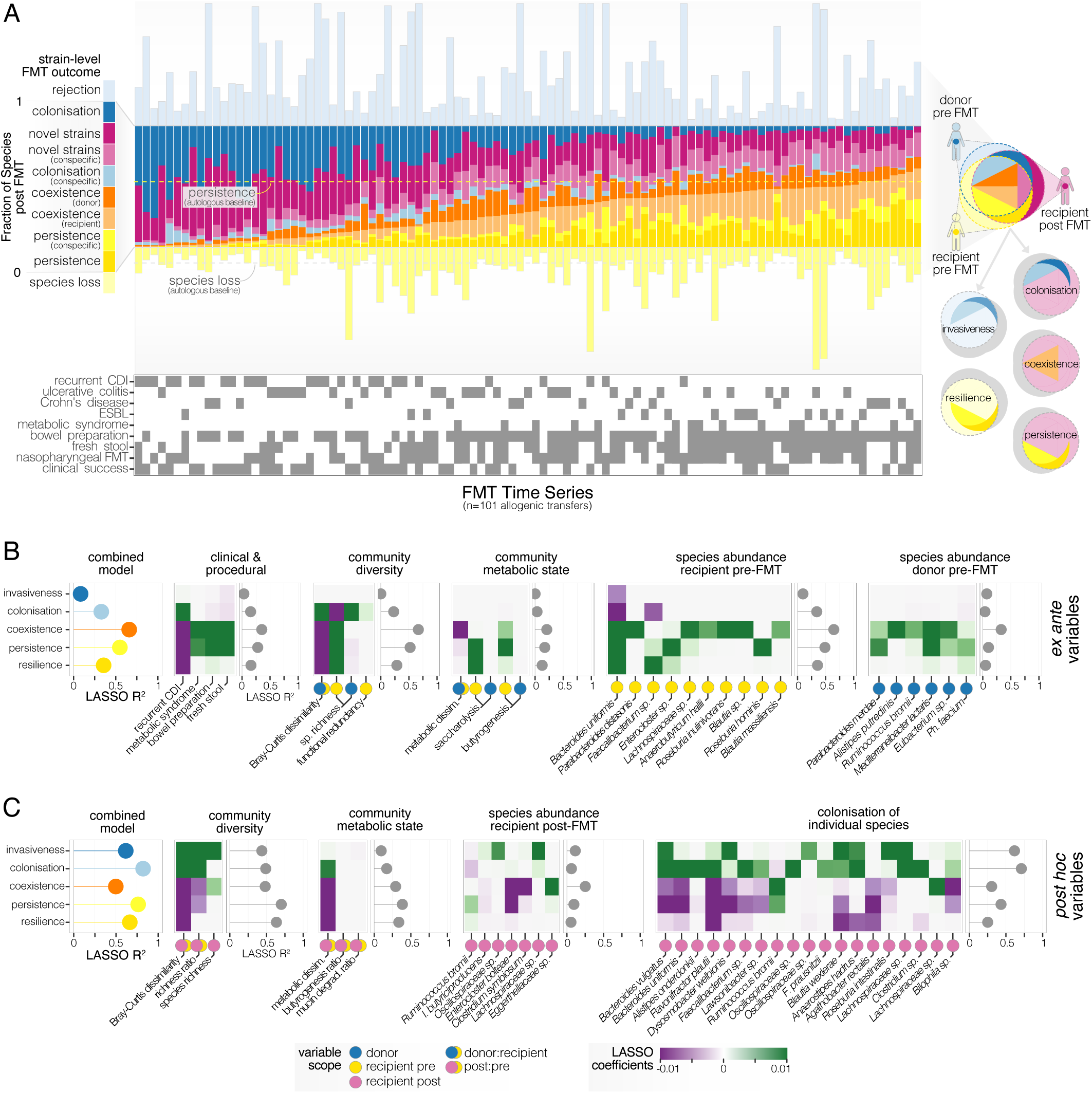
Community-wide FMT outcomes vary across patients and indications. (A) Microbiome-level outcomes of 101 scorable allogenic FMT time series, summarised across all strain populations observed in donor and recipient. Fractions are normalised to the number of species observed in the recipient post FMT, with contextual data on indication, procedure and clinical outcome. (B) *Ex ante* predictability of community-wide FMT outcomes for cross-validated LASSO linear models using regularised subsets of different variable categories or a combination of all variables that are knowable before the intervention (see Methods and table S6). Predictive performance for each outcome index is shown as R^2^, variable importance and directionality for the most predictive factors as cross-validated LASSO coefficients. (C) Association of FMT outcomes with LASSO regularised sets of *post hoc* variables (measured after the intervention).

Takeover by donor and novel strains was characteristic of rCDI and UC patients, whereas metabolic syndrome FMTs mostly resulted in conspecific strain coexistence, with intermediate outcomes in CD and ESBL. Clinical response was not associated with strain-level dynamics for any indication; in other words, patient remission was not significantly linked with donor strain colonisation or recipient strain displacement, neither for individual species nor across all tracked species (Fig S1). In particular, our data did not support earlier hypotheses that a reinstatement of short chain fatty acid (SCFA) production is a hallmark of remission in UC and CD, as an increased carriage of gut metabolic modules (GMMs, see Methods) for acetogenesis, propionigenesis and butyrogenesis following FMT did not correlate with clinical outcome. We also did not detect a notable displacement of potentially ESBL-producing *Enterobacteriaceae* (such as *E. coli* or *K. pneumoniae*) nor indeed a significant decrease in ESBL gene carriage among ESBL patients with a clinical FMT response (i.e. loss of clinically detectable ESBL). In fact, ESBL FMT responders anecdotally showed higher recipient strain persistence and less pronounced shifts in overall community composition than non-responders, indicating that the dynamics of most strain populations may be decoupled from clinical effects.

### Recipient factors outweigh donor factors when predicting FMT outcomes

To identify factors associated with FMT outcomes, we next trained a series of predictive machine learning models, using cross-validated LASSO-regularised linear regression (Methods). Among possible predictors, we distinguished *ex ante* variables (i.e. knowable prior to the FMT intervention; Fig 2B) from *post hoc* variables (measurable after FMT; Fig 2C). Moreover, we categorised predictors based on variable scope (procedural, donor-related and recipient-related) and resolution (host-, community- and species-level), totalling >400 variables as regularisation inputs (table S6). We then built cross-validated models for individual predictor categories (e.g. using procedural variables only), as well as combined models to assess overall predictability of outcomes.

Using regularised combinations of *ex ante* variables, the fraction of species exhibiting post-FMT strain co-existence (R^2^=0.65; indices as defined above, see Fig 2A) and post-FMT recipient persistence (R^2^=0.55) were predictable with moderate accuracy, with lower variance explained for colonisation by donor (R^2^=0.35) and pre-FMT recipient strain resilience (R^2^=0.35; Fig 2B). The fraction of pre-FMT donor strains to successfully take over (‘invasiveness’, R^2^=0.09) was not predictable *ex ante*. The most important outcome predictors included recipient species richness, donor-recipient community dissimilarity and recipient pre-FMT abundances of selected species, in particular *Bacteroides uniformis* (positively associated with overall strain persistence and coexistence). Recipient species richness alone was more predictive than procedural variables, such as disease indication, implying that e.g. the high levels of donor colonisation among rCDI patients may be due to an overall more disbalanced community, rather than disease-specific effects. Metabolic variables, including donor carriage of SCFA synthesis GMMs, were not predictive, although higher butyrogenesis potential in the recipient was moderately associated with overall strain persistence.

Models restricted to community diversity variables or pre-FMT recipient species abundances achieved similar accuracies to combined models choosing from all variables. In contrast, models using procedural, metabolic or donor species variables showed little or no accuracy. Across combined and individual models, variables capturing recipient factors or donor-recipient complementarity (e.g. community dissimilarity) were much more predictive than donor factors.

As expected, models trained on *post hoc* variables were generally more accurate, in particular when describing donor colonisation and invasiveness (Fig 2C). In particular, the strength of community-wide compositional shifts in the recipient (Bray-Curtis dissimilarity and metabolic dissimilarity pre- to post-FMT) were the best predictors of lower strain persistence and resilience. Interestingly, no individual species’ abundance post-FMT was associated with overall outcome. However, the carriage of individual species (Fig 2C, right) was highly predictive of overall colonisation and invasiveness, in particular *B. uniformis, B. vulgatus*, several *Oscillospiraceae sp*. including *Dysosmobacter welbionis*, and *Lachnospiraceae sp*. These might be considered indicator species, the successful engraftment of which is associated with an overall higher influx of donor strains.

### Species-specific FMT outcomes are highly variable, but predictable ex ante

Whereas the above analyses describe summarised outcomes across all tracked species, we next focussed on species-specific strain population dynamics across FMTs for 159 prevalent species, observable in ≥50% of allogenic FMTs, allowing for sufficient statistical power (Fig 3, S1 & S2). Recipient persistence, donor colonisation, coexistence and influx of novel strains were observed for all species, with no notable phylogenetic signal. No species followed consistent patterns of colonisation or persistence across all FMTs, i.e. we observed no ‘super-colonisers’ or ‘super-persisters’; rather, all species experienced all strain-level outcomes, depending on context. Strains of sporulating (ANOVA, R^2^=0.027, p=0.04) and facultatively aerobic (R^2^=0.021, p=0.005) species colonised less successfully, whereas carriage of butyrogenesis (R^2^=0.046, p=3×10^−5^) or propionigenesis (R^2^=0.02, p=0.005) pathway genes were associated with higher colonisation success. However, these mild phenotype associations were fully explained by species phylogeny and were not significant upon phylogenetic generalised least squares regression.

**Figure 3.**
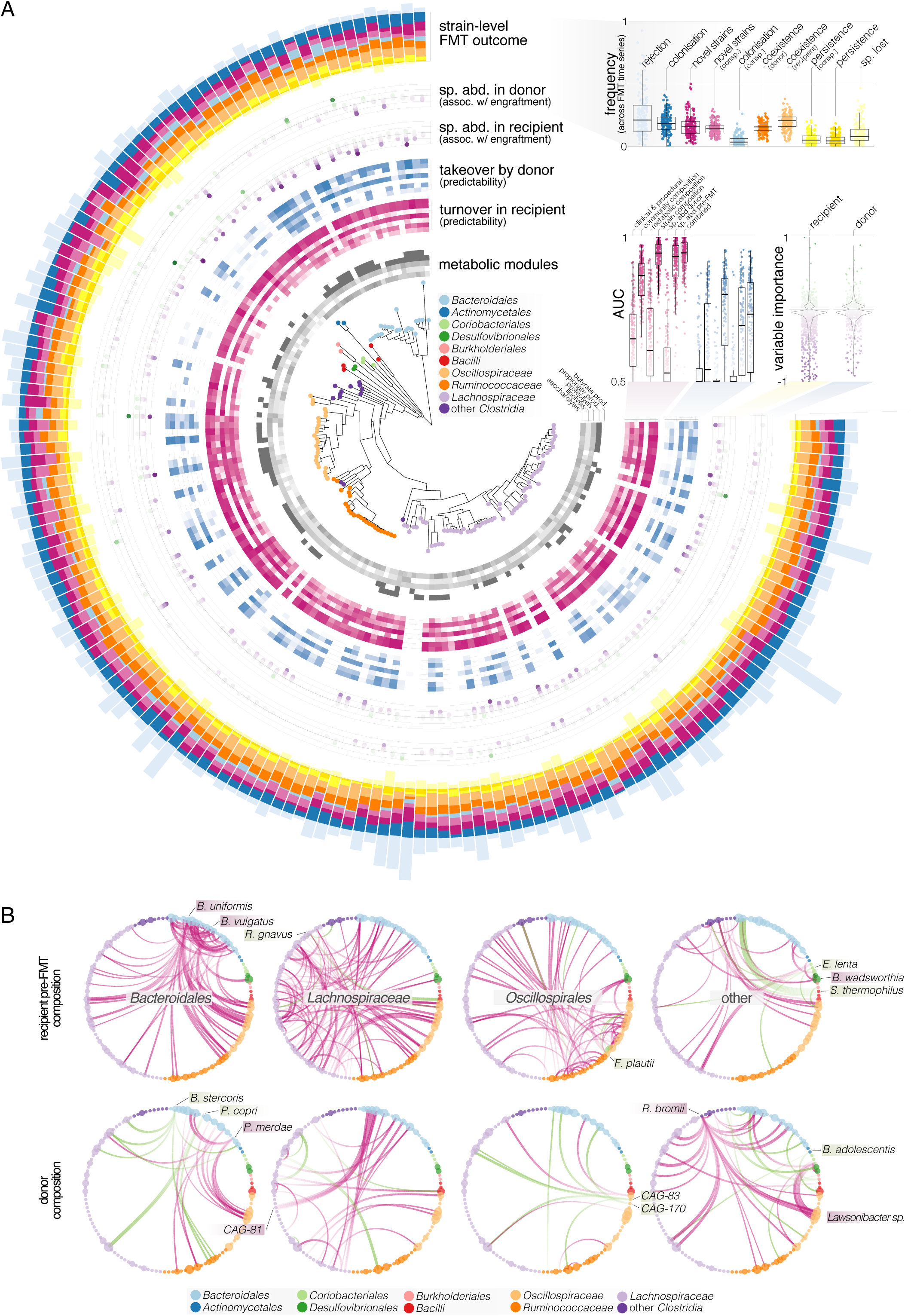
Species-level FMT outcomes depend on biotic context. (A) Phylogeny of 159 prevalent species and their strain-level outcomes across FMTs (outermost ring and summary plot top right). For each species, the predictability of binarised outcomes (recipient strain population turnover, purple ring; donor takeover, blue) using regularised subsets of *ex ante* variables are shown as AUC values, averaged across LASSO cross-validation folds. The association of each species’ abundance in donor or recipient with the successful colonisation by *other* species are shown as outer rings, summarised across species as violin plots on the right. (B) Networks of colonisation facilitation and inhibition by the recipient (top) and donor (bottom) baseline flora. Each circular subnetwork shows negative (purple edges) and positive (green) associations of members of a selected clade with the engraftment of other species. For example, high recipient pre-FMT abundances of *B. vulgatus* or *B. uniformis* correlate with lower colonisation success of several other species (top left), indicating exclusion effects. In contrast, higher donor levels of *B. stercoris* or *P. copri* among others may facilitate the colonisation by other species.

To disentangle the factors contributing to FMT outcome of each species, we built species-specific cross-validated logistic LASSO regression models, using *ex ante* and *post hoc* predictor sets analogously to those discussed above. We binarised species outcomes, defining *donor takeover* events as all outcomes with dominant colonisation by donor strains, *recipient turnover* as outcomes with a displacement of recipient strains (i.e. species loss, donor colonisation or influx of novel strains), and *recipient resilience* as outcomes where recipient strains persisted as dominant or coexisting populations. The predictability of both takeover (blue ring in Fig 3A) and turnover (purple) based on *ex ante* variables varied across clades: whereas outcomes for *Bacteroidales sp*. were predictable with high accuracy (Area Under the Receiver Operator Characteristic curve, AUC≥0.85; small inset in Fig 3A), predictive performance was close to random for several clades of Actinobacteria and Clostridia. In general, recipient strain turnover (AUC for full models, 0.94±0.06) was more predictable than donor takeover (AUC=0.70±0.15) and recipient resilience (AUC=0.57±0.12) across all species, indicating that the displacement of native recipient strain populations by any means (not just replacement by donor strains) may be a more deterministic process in general.

These signals were driven by specific subsets of predictive variables (Fig 4A). Models relying exclusively on *ex ante* microbiome community composition, pre-FMT species abundances in the recipient or strain composition of the focal species achieved highest accuracies, comparable to those of full combined models, whereas procedural factors, metabolic and donor species abundances were much less predictive.

**Figure 4.**
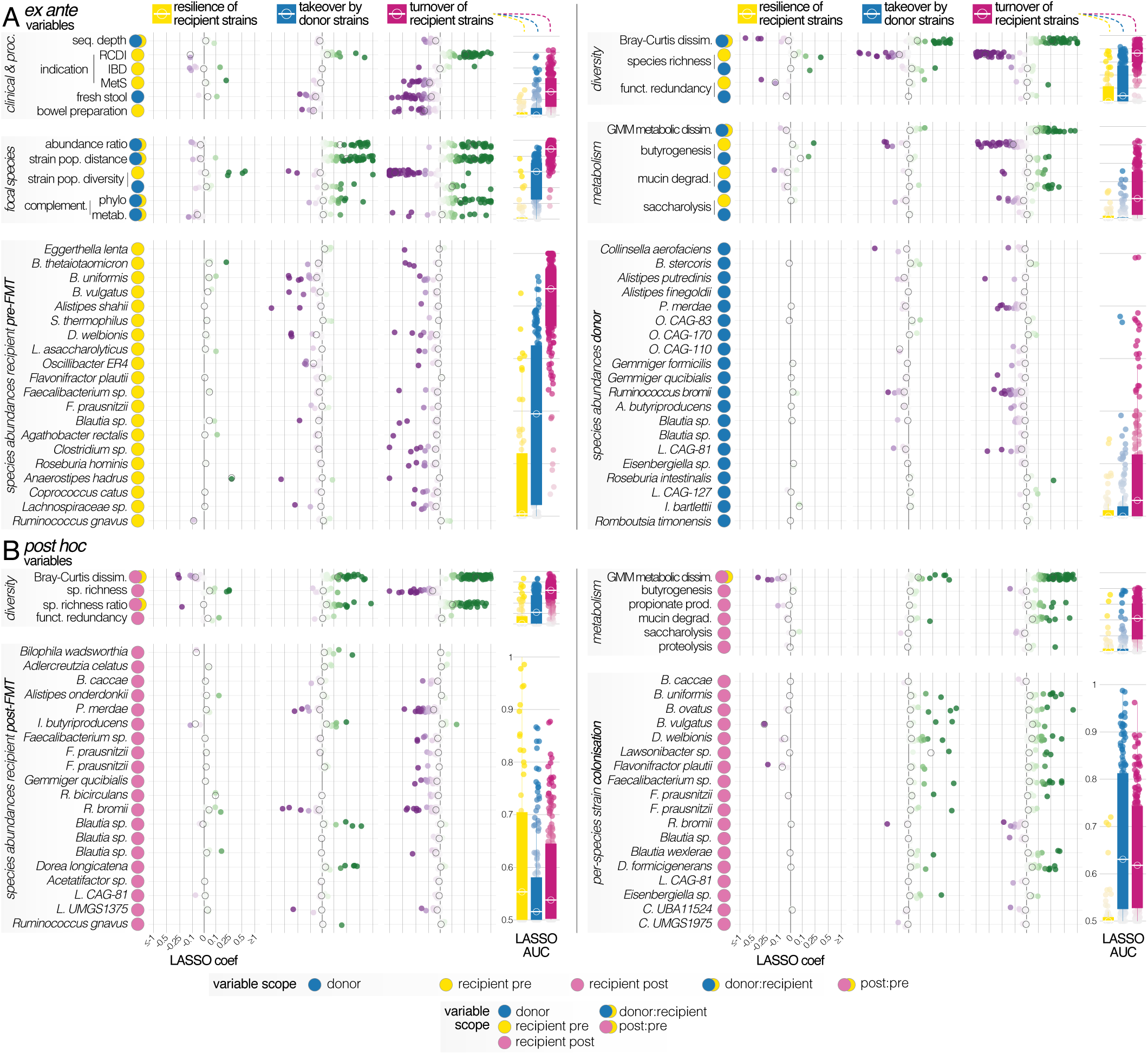
Predictability of recipient strain population resilience, turnover and donor takeover for individual species. (A) Logistic LASSO models were trained to predict FMT binarised outcomes (recipient resilience, yellow; recipient turnover, purple; donor takeover, blue) per species across FMT time series, using different subsets of *ex ante* variables (knowable before the intervention). Each dot represents data for one species, variables are categorised by scope (donor, recipient) and type (procedural, community-level diversity, etc.). Predictive performance of species models is shown as average AUC across LASSO cross-validation folds in marginal box plots, ranging from [0.5, 1]. Variable importance within each category and association directionality are shown as averaged standardised LASSO coefficients; median values across species are indicated as void circles. The bottom panels (species abundances in donor/recipient) correspond to subsets of the networks shown in Fig 3B. (B) Association of *post hoc* variables (measured after the intervention) with FMT outcomes for individual species. The successful colonisation by several ‘indicator species’ was moderately associated with FMT outcome for other, unrelated species (bottom right panel).

### Strain turnover is driven by recipient community state and donor-recipient complementarity

The use of only two coarse-grained microbiome diversity variables – recipient species richness pre-FMT and donor-recipient community dissimilarity – were sufficient to accurately predict strain turnover for most species (Fig 4A). Low richness and a strong compositional shift in the recipient microbiome relative to healthy donors are hallmarks of disease-associated microbiome states, and our data indicates that the strength of this diffuse disbalance, correlated to disease (such as rCDI or UC in our dataset), is directly linked with FMT outcome in most species. In contrast, donor richness or functional redundancy, previously proposed to be relevant ^43^, were only subordinately predictive, if at all. Metabolic variables were likewise unreliable predictors. Community-wide butyrogenesis potential was negatively associated with turnover in the recipient (i.e. strain populations were more resilient in recipients carrying high loads of butyrate production genes), but higher butyrogenesis levels in the donor did not correspondingly promote colonisation.

For each species, strain-level and functional properties, and in particular the complementarity between donor and recipient populations, were excellent predictors of FMT colonisation outcome. The donor-recipient abundance ratio was also a relevant predictor, as suggested previously ^27^. This indicates that mere propagule pressure (the amount of incoming viable donor microbes) may provide a neutral baseline estimate for colonisation success, in particular for species not present in the recipient pre-FMT. However, infra-specific effects were indeed more subtle than that: donor-recipient strain population dissimilarity and recipient (and to a lesser extent, donor) strain population diversity were superior outcome predictors. In other words, turnover or colonisation were more likely in species with *complementary* strain populations between donor and recipient, while diverse recipient populations (not dominated by individual strains) were more resilient than uneven ones. Moreover, incoming species that were phylogenetically or metabolically complementary to the recipient community (i.e. adding novelty, e.g. by filling an unoccupied niche) were more likely to colonise or turn over the resident population.

We observed similar trends for *post hoc* variables (Fig 4B). Interestingly, some species were decent engraftment or turnover indicators: for example, the successful engraftment of several *Bacteroides* sp., *Faecalibacterium* sp., *Dysosmobacter welbonis* or *Dorea formicigenerans* were indicative of outcomes for other species, albeit to a limited extent.

### Resident flora of the recipient negatively impacts donor colonisation, with notable exceptions

Given that FMTs involve the pitting of the resident’s residual microbial community against incoming flora from the donor, we explored the specific impact of individual species on the engraftment of others (Fig 3B and 4A). We built networks of engraftment inhibition and facilitation, associating the abundance of putative effector species in the donor and recipient with donor takeover events in focal species. The vast majority of interactions was inhibitive (outer rings and inset in Fig 3A): for most species, higher residual abundance in both donor and recipient correlated negatively with engraftment of other species. Recipient abundances (AUC=0.68±0.15) were more informative than donor abundances which were hardly predictive of colonisation at all (AUC=0.52±0.06).

Colonisation inhibition was phylogenetically concentrated, i.e. inhibitive interactions were more common between related species within the same clade than between clades (Fig 3B). *Bacteroides* sp. in the recipient flora, in particular *B. uniformis* and *B. vulgatus*, were among the strongest colonisation inhibitors, but also included the most strongly inhibited species, *B. xylanisolvens* and *B. ovatus*. In other words, the enrichment of several *Bacteroides* sp. in the recipient microbiota inhibited colonisation for a broad panel of species, and vice versa, in line with previous findings that *Bacteroides* are generally highly persistent also in healthy individuals^44^. *Ruminococcus gnavus, Eggerthella lenta, Flavonifractor plautii* and *Streptococcus thermophilus* in the recipient were the foremost colonisation facilitators. In contrast to colonisation inhibition, facilitation typically affected phylogenetically distant species: for example, the facilitation of *B. stercoris* colonisation by recipient *S. thermophilus* was the strongest interaction observed across all species.

We observed a few prominent colonisation-facilitating species in the donor flora, most notably *B. stercoris, Prevotella copri* and *Bifidobacterium adolescentis*. However, facilitation and inhibition effects of donor species were generally limited and overall less predictive of colonisation success, indicating that the donor flora has small impact on FMT outcome, beyond intra-specific strain dynamics.

### Both neutral and adaptive processes shape post-FMT strain population dynamics

The accurate prediction of FMT outcomes is informative beyond mere descriptive associations when construed through the lens of gut ecology: FMTs are perturbation experiments *in natura*, interpretable in a framework of invasion ecology and community assembly to identify processes and mechanisms that shape the microbiome ^33,34^. We therefore linked the various tested variables in our models to putative underlying mechanisms (Fig 5), categorised along a gradient from neutral/stochastic factors (e.g. donor propagule pressure) to adaptive/selective ones (e.g. niche effects). We further distinguished recipient-specific, donor-specific and donor-recipient complementarity effects, and organised variables by granularity, from host-level factors (e.g. clinical or procedural) to the level of microbiome communities (overall composition and possible species interactions) and intra-specific (strain-level) effects.

**Figure 5.**
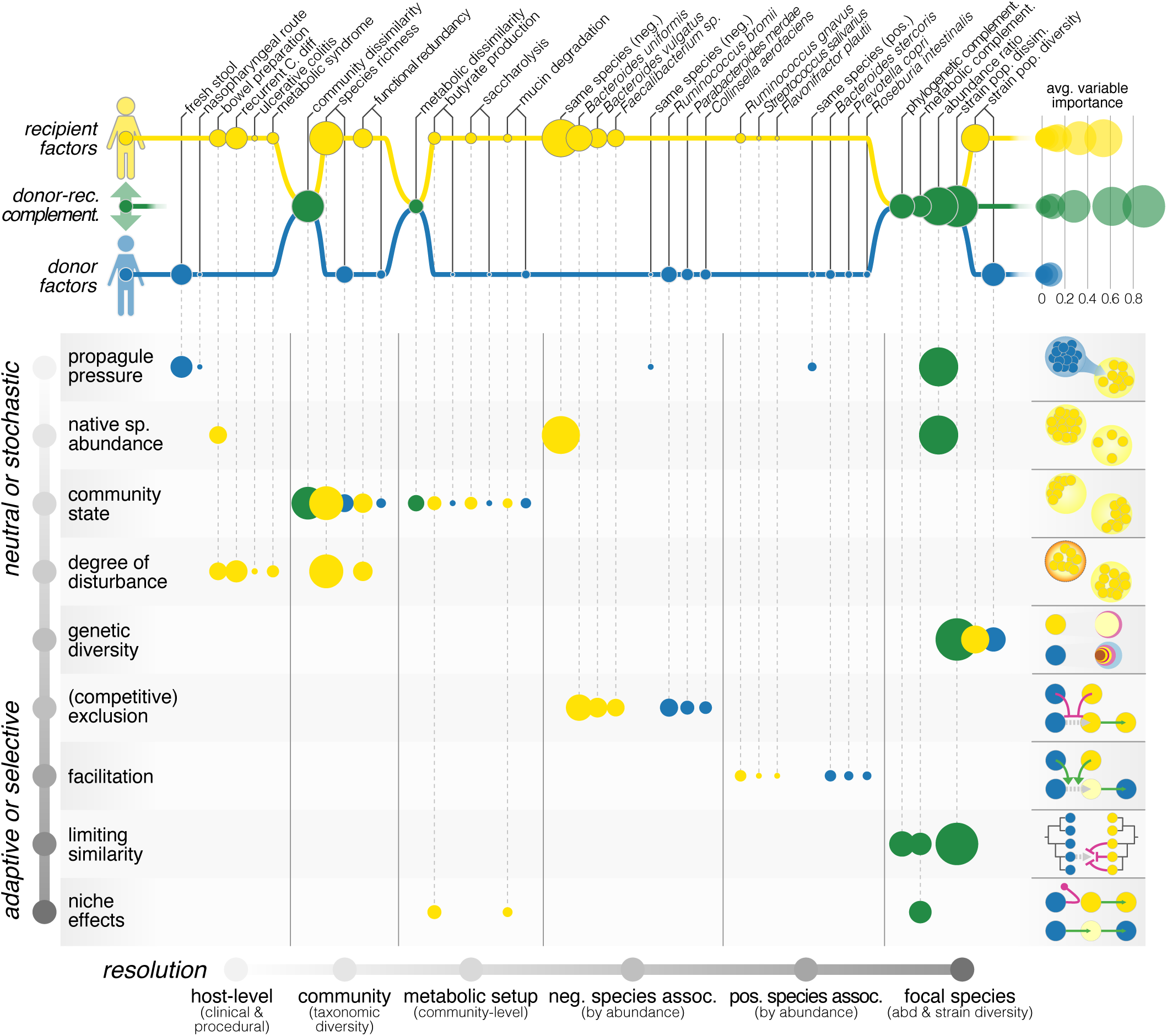
FMT outcomes are shaped by both neutral and adaptive ecological processes. Each of the tested variables to predict FMT outcome can be linked to putative underlying ecological processes, as suggested previously ^33^. Factors are organised by scope (pertaining to the donor, recipient, or donor-recipient complementarity; top) and resolution (host-, community-, species- and strain-level; left to right). Underlying ecological processes can be roughly ranked along the gradient from neutral/stochastic to adaptive/selective; each process is illustrated with a toy example on the right. Circle size corresponds to average variable importance, calculated across all tested species from LASSO coefficients and overall model performance (less predictive models penalise variable importance). Recipient factors, and in particular donor-recipient complementarity measures across all resolutions were generally far more relevant to species-level outcome than donor factors (top right sub-panel).

Factors pertaining to the recipient or to donor-recipient complementarity were far more relevant to FMT outcome than donor readouts across all tested variables, and consistently across different species. In other words, the donor microbiome did not specifically influence colonisation or turnover in its own right, but instead mattered only to the extent of its complementarity with the recipient microbiota. Donor-recipient abundance ratios were highly determinant of FMT outcome, interpretable as the balance between propagule pressure of incoming donor cells and native abundance of the residual recipient population, providing a baseline of how neutral mechanisms shape post-FMT communities. In this, exclusion effects by native populations were dominant, i.e. higher donor propagule dosage was less relevant to successful colonisation than a depletion of the residual microbiota. In practice, this interplay may be modulated procedurally to some extent, e.g. by the use of fresh versus frozen stool (impacting the viability of donor cells), FMT route (rectal or duodenal), or the purging of recipient communities via bowel preparations or antibiotics pre-treatment.

General microbiome compositional state in the recipient (but not the donor) was likewise relevant to FMT outcome: broad community depletion (low richness) and pronounced compositional differences from healthy donors may indicate generally disturbed and precarious microbiome states, less resistant to donor takeover. Conversely, the residual enrichment of ‘gatekeeper’ species such as *B. uniformis* or *B. vulgatus* was also negatively associated with donor colonisation, possibly indicating competitive exclusion processes and inter-specific priority effects. While by design, causality cannot be inferred from our data, these results tie in with existing ecological theories on microbiome stability and resilience, e.g. on tipping elements and critical transitions ^45,46^, community multi-stability leading to enterotypes ^47,48^, priority ^49^ or Anna Karenina effects ^50^. We found limited evidence for colonisation facilitation across species boundaries, both in donor and recipient. Likewise, our data did not support a strong role for community-wide metabolic states: neither general metabolic setup nor specific metabolic modules such as SCFA production in donor or recipient greatly impacted FMT outcomes.

The strongest effects toward FMT outcome emerged at species and strain level. Incoming species were more likely to colonise if they were phylogenetically or metabolically complementary to the residual community, implying that they were able to take over unoccupied niches. Colonisation success was associated with complementarity specifically to the local community. High intraspecific diversity in the donor and low diversity in the recipient were also linked with engraftment success: recipient populations dominated by single strains were less resilient, and donor strains from more diverse panels were more invasive, likely due to strain-level limiting similarity effects. Indeed, conspecific donor strain populations colonised more successfully if they were dissimilar to recipient strains, indicating strong inhibitive intra-specific priority effects.

However, we note once more that the colonisation of individual species was only predictable with moderate accuracy, irrespective of the variable sets used – unlike residual strain population turnover which was highly predictable. This implies that colonisation success may be stochastic to a large extent.

## Discussion

Faecal microbiota transplantations are usually conducted for clinical purposes, but can also be thought of as complex *in natura* perturbation experiments, pitting donor against recipient microbial communities. From a clinical perspective, an FMT is successful if it triggers patient remission or recovery, whereas success from an ecological perspective is the extent to which the donor’s flora can colonise the recipient. Given that FMT targets the microbiome, engraftment and clinical success are expected to correlate, implying that successful microbiome modulation mediates clinical effects. However, this hypothesis had previously not been systematically tested, and is indeed not supported by our data. In our meta-study of 142 FMTs, clinical success was neither associated with donor colonisation, the displacement of recipient species nor the reinstatement of specific functions (such as SCFA synthesis) for any of the five studied indications. To some extent, this is in line with previous observations that autologous FMTs ^51,52^ or even transfers of sterile-filtered faecal water ^53^ can be efficacious. Our data does not rule out more subtle links, in particular given our limited sample size per indication, but a clear role of donor microbiota colonisation in shaping clinical responses did not emerge. We did observe overall higher levels of donor colonisation in patients suffering from rCDI or UC, coinciding with higher clinical response rates in these diseases compared to others in our dataset. However, this was arguably due to overall more perturbed microbiome states associated with these diseases that outweighed disease-specific effects: we found no significant differences in strain-level outcomes between FMT responders and non-responders.

Understanding microbiome-level FMT outcomes is both clinically relevant (e.g. for informed donor selection or to avoid possible adverse effects) and more generally informative of ecological processes shaping the gut microbiome. All studied species exhibited all FMT outcomes, depending on context; we did not find strong evidence that any species were inherently more invasive or resilient than others. Rather, fine-scale intra-specific strain population structure and diversity, as well as donor-recipient strain population complementarity determined resilience, coexistence and colonisation. Interactions between species were less relevant, but clearly structured: several ‘gatekeeper’ species in the recipient, in particular of the genus *Bacteroides*, inhibited colonisation by other, phylogenetically unrelated species, whereas colonisation facilitation across species boundaries was scarce.

We found that the displacement (turnover) of recipient strains was very accurately predictable for almost all studied species, using a consistent and surprisingly small selection of *ex ante* microbiome variables. In contrast, our models achieved only moderate predictive accuracies for donor strain takeover, indicating that colonisation is to a large extent stochastic, or influenced by other factors outside the scope of our study, such as viral or eukaryotic microbiome members, recipient immune state or reduced viability of anaerobic donor faecal cells following the intervention.

Recipient factors consistently outweighed donor factors in driving FMT outcomes. Thus, our data did not support the ‘super donor’ hypothesis ^17^ which states that certain donor microbiome properties are crucial to colonisation and, by proxy, clinical success. Neither did we observe any consistent effects of using relatives or cohabitants as donors. Rather, we found that donor-recipient *complementarity* promoted donor colonisation and recipient turnover. Complementarity mattered across resolutions, from community-level effects to intra-specific strain population dissimilarity. Indeed, strain-level diversity and complementarity were the strongest determinants of FMT outcome, with relevance to rational donor selection in the clinical practice ^18^. Beyond screening for donor health, matching donors to recipients based on microbiome complementarity at community, species and in particular strain levels may increase colonisation success, make outcomes more predictable, and reduce adverse effects.

Our data suggests that the gut microbiome is shaped by both neutral and adaptive processes post FMT, reconciling previous anecdotal reports ^27,32^. We found that limits to gut microbiome resilience at community, species and strain level can be defined by a relatively small set of measurable variables that point to distinct underlying processes. The (complementary) interplay between propagule pressure and residual species abundance provided a neutral baseline for colonisation, although again, recipient outweighed donor effects. At the same time, our data also suggested niche effects, in particular at the level of complementary intra-specific strain populations, although no consistently adaptive traits emerged in the analysis. Hypotheses pertaining to the importance of metabolic capabilities such as SCFA synthesis were not supported, although we note that the inference of SCFA biosynthesis pathways from metagenomic data remains challenging and does not capture putatively differential expression of SCFA synthesis genes.

Our findings may inform the clinical use of FMT in several ways, in particular if microbiome modulation is a desired endpoint beyond alleviation or remission of symptoms. Patients may be stratified prior to the intervention based on surprisingly crude, robust and easily obtainable microbiome readouts, such as community richness and high-level composition, or with regard to the presence of ‘gatekeeper’ species associated with overall microbiome resilience. The relevance of donor selection, in contrast, appears mostly limited to the extent of the donor’s (strain-level) complementarity to the recipient. Tuning procedural parameters (stool preparation, dosage, FMT route, dietary intake of donors, etc.) may mostly impact recipient microbiome resilience, and an overall more resilient response (excluding of course target pathogens to be displaced) is often desirable. Both inhibition and facilitation of outcomes across species boundaries were surprisingly sparse and mild, indicating that the targeted colonisation or turnover of individual species may be achievable mostly independent of residual and co-transferred communities, minimising collateral effects on the recipient’s flora. Our findings strongly support the notion that predictable and efficacious microbiome modulation using personalised probiotic mixtures, rather than entire complex faecal samples, is possible and will greatly profit from an ecological perspective. In particular, our data implies that the concurrent use of diverse intra-specific strain populations, rather than individual strains, may greatly enhance colonisation and turnover in the recipient. Thus, levering of both neutral and relevant adaptive ecological processes may pave the way towards targeted modulatory interventions on the gut microbiome, personalised to recipients, and with predictable microbiome-level outcomes.

## Methods

### Data overview

The study dataset comprised of 12 independent cohorts, recruited in centers in the USA, the Netherlands and Australia, with a total of 142 FMTs conducted in 134 patients suffering from recurrent *Clostridioides difficile* infection (rCDI, n=37 ^26,28,38^), ulcerative colitis (UC, n=42 ^30,35–37^), Crohn’s disease (CD, n=18 ^41^), metabolic syndrome (MetS, n=26 ^25,39^) or infection with extended-spectrum beta lactamase producing bacteria (ESBL, n=19 ^40^). On average, 3.26 recipient stool samples were available per FMT time series, including baseline samples taken prior to the intervention (‘pre-FMT’). Overall, 3.5 Tbp of sequencing data were analysed across 556 faecal metagenomes, of which 269 (for 76 time series) were generated as part of the present study (for cohorts NL_UC, NL_ESBL, NL_MetS and AU_div).

Two cohorts (NL_UC and NL_MetS) were randomised controlled trials during which a subset of patients received autologous FMTs (transplantation of the recipient’s own stool, n=21). All other FMTs were allogenic, using stool donors. For 122 FMT time series, a full complement of donor baseline, recipient baseline and at least one recipient post-FMT sample were available.

A full description of all cohorts is provided in table S1; detailed information per FMT time series in table S2; and per-sample information in table S3.

### Sample collection, processing and metagenomic sequencing

Study design and faecal sample collection for the cohorts NL_MetS ^25,39^, NL_UC ^35,54^ and NL_ESBL ^40^ were described previously. AU_rCDI and AU_UC samples were obtained from a single-centre, proof-of-concept, parallel and controlled study in collaboration with the Centre for Digestive Diseases (Sydney, Australia), which aimed to assess donor microbiota implantation in 2 CDI and 3 UC patients up to 28 days following a 2-day faecal microbiota transplantation infusion via trans-colonoscopy and rectal enema.

For the NL_MetS and NL_UC cohorts, faecal DNA extraction was described in the original studies. DNA from NL_ESBL samples was extracted using the GNOME® DNA Isolation Kit (MP Biomedicals) with the following minor modifications: cell lysis/denaturation was performed (30 min, 55°C) before protease digestion was carried out overnight (55°C), and RNAse digestion (50 μl, 30 min, 55°C) was performed after mechanical lysis. After final precipitation, the DNA was resuspended in TE buffer and stored at −20°C for further analysis.

Metagenomic sequencing libraries for NL_MetS, NL_UC, NL_ESBL, AU_rCDI and AU_UC samples were prepared to a target insert size of 350-400bp on a Biomek FXp Dual Hybrid, with high-density layout adaptors, orbital shaker, static peltier, shaking peltier (Beckman Coulter, Brea, USA), and a robotic PCR cycler (Biometra, Göttingen, Germany), using SPRIworks HT kits (Beckman Coulter) according to the supplier’s recommendation with the following modifications: 500 ng DNA starting amount, adaptor dilution 1:25, kit chemical dilution 1:1 in process. For samples with low input DNA concentrations, libraries were instead prepared manually using NEBNext Ultra II DNA Library Prep kits with NEBNext Singleplex primers. Libraries were sequenced on an Illumina HiSeq 4000 platform (Illumina, San Diego, CA, USA) with 2×150bp paired-end reads.

### Public datasets

Based on a literature search, 8 datasets on FMT cohorts that met the following criteria were included in the study: (i) public availability of metagenomic sequencing data in September 2018; (ii) sufficient available description to unambiguously match donors and recipients per FMT time series; (iii) no restrictions on data reuse. They were included in this study as US_rCDI_1 (n=22 FMT time series ^26^), US_rCDI_2 (n=7 ^28^), US_rCDI_3 (n=6 ^38^), US_UC_1 (n=6 ^37^), US_UC_2 (n=4 ^36^), US_UC_3 (n=2 ^30^), and US_CD (n=18 ^41^). Contextual data, including donor-recipient matchings or information about clinical response, were curated from the study publications and in some cases kindly amended by the studies’ original authors upon request (see tables S1-S3).

### Metagenomic data processing, taxonomic and functional profiling

Metagenomic reads were quality trimmed to remove base calls with a Phred score of less than 25. Reads were then discarded if they were shorter than 45 nucleotides or if they mapped to the human genome (GRCh38.p10) with at least 90% identity over 45 nucleotides. This processing was performed using NGLess ^55^. Taxonomic profiles per sample were obtained using mOTUs v2 ^56^. For functional profiling, reads were mapped against the Global Microbial Gene Catalogue v1 gut sub-catalogue (gmgc.embl.de, Coelho et al. *in revision*) with a minimum match length of 45 nucleotides with at least 97% identity and summarised based on antimicrobial resistance gene (ARG) annotations and KEGG orthologs (KOs) via eggNOG annotations ^57^. Based on the resulting KO profiles, Gut Metabolic Modules (GMMs ^58^) were quantified in each sample using omixer-rpmR ^59^. Taxonomic and GMM profiles per sample, normalised by read depth, are available as supplementary tables S7 and S8.

### Metagenome-Assembled Genomes (MAGs)

We demarcated Metagenome-Assembled Genomes (MAGs) using several complementary strategies to obtain both high resolution from sample-specific assemblies and deep coverage of lowly abundant species from co-assemblies of multiple samples. Unless otherwise indicated, all tools in the following were run with default parameters.

To generate single-sample MAGs, faecal metagenomes were assembled individually using metaSPAdes v3.12.0 ^60^, reads were mapped back to contigs using bwa-mem v0.7.17 ^61^, and contigs were binned using metaBAT v2.12.1 ^62^. Multi-sample MAGs were built for each cohort separately. Reads were first co-assembled using megahit v1.1.3 ^63^ and mapped back to contigs using bwa-mem v0.7.17. Co-assembled contigs were then binned using both CONCOCT v0.5.0 ^64^ and metaBAT v2.12.1. The resulting co-assembled MAG sets were further refined using DAS TOOL ^65^ and metaWRAP ^66^. In total, 47,548 MAGs were demarcated using these five approaches (single-sample MAGs, multi-sample co-assembled CONCOCT, metaBAT2, DAS TOOL and metaWRAP MAGs). In addition, we included 25,037 high quality reference genomes from the proGenomes database ^42,67^ in downstream analyses.

Genome quality was estimated using CheckM ^68^ and GUNC v0.1 ^69^, and all genomes were taxonomically classified using GTDB-tk ^70^. Open reading frames (ORFs) were predicted using prodigal ^71^ and annotated via the prokka workflow v1.14.6 ^72^. Orthologs to known gene families were detected using eggNOG-mapper v1 ^73^. ARGs were annotated using a workflow combining information from the CARD v3.0.0 (via rgi v4.2.4, ^74^) and ResFams v1.2.2 ^75^ databases, as described previously ^67^. The ‘specI’ set of 40 near-universal single copy marker genes were detected in each genome using fetchMG ^76^.

The full set of generated MAGs and contextual data is available via Zenodo.

### Genome clustering, species meta-pangenomes, phylogeny

Genomes were clustered into species-level groups using an ‘open-reference’ approach in multiple steps. Initial pre-filtering using lenient quality criteria (CheckM-estimated completeness ≥70%, contamination ≤25%; additional criteria were applied downstream) removed 57.7% of MAGs. The remaining 20,093 MAGs were mapped to the clustered proGenomes v1 ^42^ and mOTUs v2 ^56^ taxonomic marker gene databases using MAPseq v1.2.3 ^77^. 17,720 MAGs were confidently assigned to a ref-mOTU (specI cluster) or meta-mOTU based on the following criteria: (i) detection of at least 20% of the screened taxonomic marker genes and (ii) a majority of markers assigning to the same mOTU at a conservative MAPseq confidence threshold of ≥0.9.

In an independent approach, quality-filtered MAGs and reference genomes were also clustered by average nucleotide identity (ANI), using a modified and scalable reimplementation of the dRep workflow ^78^. Using pairwise distances computed with mash v2.1 ^79^, sequences were first pre-clustered to 90% mash-ANI using the single linkage algorithm, asserting that all genome pairs sharing ≥90% were grouped together. Each mash pre-cluster was then resolved to 95% and 99% average linkage ANI clusters using fastANI v1.1 ^80^. For each cluster, a representative genome was picked as either the corresponding reference specI cluster representative in the proGenomes database, or the MAG with the highest dRep score (calculated based on estimated completeness and contamination). Genome partitions based on 95% average linkage ANI clustering and specI marker gene mappings matched almost perfectly, at an Adjusted Rand Index of >0.99. We therefore defined a total of 1,056 species-level clusters (‘species’) from our dataset (see table S4), primarily based on marker gene mappings to pre-computed ref-mOTUs (or ‘specI’ clusters, n=295) and meta-mOTUs (n=528), and as 95% average linkage ANI clusters for genomes that did not map to either of these databases (n=233).

Species pangenomes were generated by clustering all genes within each species-level cluster at 95% amino acid identity, using Roary 3.12.0 ^81^. Spurious and putatively contaminant gene clusters (as introduced by mis-binned contigs in MAGs) were removed by asserting that the underlying gene sequences originated (i) from a reference genome in the proGenomes database, or (ii) from at least two independent MAGs, assembled from distinct samples or studies. To account for incomplete genomes, ‘extended core genes’ were defined as gene clusters present in >80% of genomes in a species-level cluster. If too few gene clusters satisfied this criterion, as was the case for some pangenomes containing many incomplete MAGs, the 50 most prevalent gene clusters were used instead. Representative sequences for each gene cluster were picked as ORFs originating from specI representative genomes (i.e. high quality reference genomes), or else as the longest ORF in the cluster.

A phylogenetic tree of species-level cluster representatives was inferred based on the ‘mOTU’ set of 10 near-universal marker genes ^56^. Marker genes were aligned in amino acid sequence space across all species using Muscle (^82^ v 3.8.31), concatenated and then used to construct a species tree using FastTree2 (v2.1.11) ^83^ with default parameters.

### Inference of microbial strain populations

Metagenomic reads for each sample were mapped against gene cluster representative sequences for all species pangenomes using bwa-mem v0.7.17 ^61^. Mapped reads were filtered for matches of ≥45bp and ≥97% sequence identity, sorted, and filtered against multiple mappings using samtools v1.7 ^84^. Horizontal (‘breadth’) and vertical (‘depth’) coverage of each gene cluster in each sample were calculated using bedtools v2.27.1 ^85^.

A species was considered present in a sample if at least three mOTU taxonomic marker genes were confidently detected either via the mOTU v2 profiler (for specI clusters and meta-mOTUs) or based on pangenome-wide read mappings (for non-mOTU species-level clusters). Gene clusters within each pangenome were considered present in a sample if (i) the species was detectable (see previous), (ii) horizontal coverage exceeded 100bp and 20% of the representative gene’s length, and (iii) average vertical coverage exceeded 0.5. Gene clusters were considered *confidently absent* if they did not attract any mappings in samples where the species’ set of extended core genes (see above) was covered at >1 median vertical coverage (i.e. present with high confidence). Using these criteria, strain population-specific gene content profiles were computed for each species in each sample.

Raw microbial single nucleotide variants (SNVs) were called from uniquely mapping reads using metaSNV v1.0.3 ^86^ with permissive parameters (*-c 10 -t 2 -p 0*.*001 -d 1000000*). Candidate SNVs were retained if they were supported by ≥2 reads each in ≥2 samples in which the focal gene cluster was confidently detected (see above), prior to differential downstream filtering. At multi-allelic positions, the frequency of each observed allele (A, C, G, T) was normalised by the total read depth for all alleles.

Based on this data, strain populations were represented based on both their specific gene content profile and SNV profile in each sample.

Each species’ local strain population diversity (SPD) and allele distances (AD) between strain populations across samples were estimated as follows. SPD was calculated based on the inverse Simpson index of allele frequencies *p*_*[ACGT]*_ at each variant position *i* in the extended core genome, normalised by total horizontal coverage (number of covered positions) *cov*_*hor*_:

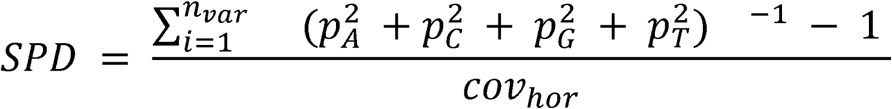

Thus defined, SPD can be interpreted as the *average effective number of non-dominant alleles* in a strain population. SPD ranges between 0 (only one dominant strain detected, i.e. no multi-allelic positions) and 3 (all four possible alleles present at equal proportions at each variant position). The normalization by total horizontal coverage *cov*_*hor*_ of the extended core genome ensures that values are comparable between samples even if a species’ coverage in a sample is incomplete.

Intra-specific allele frequency distances (AD) between strain populations across samples were calculated as the average Euclidean distance between observed allele frequencies at variant positions in the species’ extended core genome, requiring at least 20 variant positions with shared coverage between samples. If a species was not observed in a sample, AD to that sample were set to 1.

### Quantification of strain-level outcomes

Colonisation by donor strains, persistence of recipient strains and influx of ‘novel’ strains (environmental or previously below detection limit) in the recipient microbiome following FMT were quantified for every species based on determinant microbial SNVs and gene content profiles, using an approach extending previous work ^25,87^. 122 FMT time series (101 allogenic and 21 autologous transfers) for which a donor baseline (in allogenic FMTs; ‘D’), a recipient pre-FMT baseline (‘R’) and at least one recipient post-FMT (‘P’) sample were available were taken into account, and each FMT was represented as a D-R-P sample triplet. If available, multiple timepoints post FMT were scored independently. By definition, no donor samples were available for autologous FMTs, so recipient pre-FMT samples were used instead. An overview of possible strain-level FMT outcomes is provided in Figure 1C & D.

For each D-R-P sample triplet, conspecific strain dynamics were calculated if a species was observed in all three samples (see above) with at least 100 informative (‘determinant’) variant positions either covered with ≥2 reads or confidently absent (see below). Donor determinant alleles were defined as variants unique to the donor (D) relative to the recipient pre-FMT (R) sample, and vice versa. Post-FMT determinant alleles were defined as variants unique in P relative to both D and R. Given that intra-specific faecal strain populations are often heterogeneous, i.e. consist of more than one strain per species, multiple observed alleles at the same variant position were taken into account. In addition, if a gene containing a putative variant position was absent from a sample although the species’ extended core genome was detected, the variant was considered ‘confidently absent’ and treated as informative (and potentially determinant) as well, thereby taking into account differential gene content between strains.

The fraction of donor and recipient strains post FMT were quantified based on the detection of donor-determinant and recipient-determinant variants across all informative positions in the P sample. The fraction of ‘novel’ strains (environmental or previously below detection limit in donor and recipient) were quantified as the fraction of post-FMT determinant variants. Based on these three readouts (fraction of donor, recipient & ‘novel’ strains), FMT outcomes were scored categorically as ‘donor colonisation’, ‘recipient persistence’, ‘donor-recipient coexistence’ or ‘influx of novel strains’ for every species (see table S5).

In addition to conspecific strain dynamics (i.e. where a species was present in D, R & P), we also quantified FMT outcomes that involved the acquisition or loss of entire strain populations.

For example, if a species was present in the recipient at baseline, but not post-FMT, this was considered a ‘species loss’ event. See Fig 1C and table S5 for a full overview of how different FMT outcome scenarios were scored.

To assert the accuracy of our approach, we simulated FMT time series by shuffling (i) the donor sample, (ii) the recipient pre-FMT sample, or (iii) both. Randomizations were stratified by subject (accounting for the fact that some donors were used in multiple FMTs, and that some recipients received repeated treatments) and geography. For each observed D-R-P sample triplet, we simulated 10 triplets per each of the above setups.

Outcomes were further summarised across species by calculating a series of strain population-level metrics for each FMT, defined as follows (see Fig 1E for visualised definitions).

*Persistence index*: average fraction of persistent recipient strains among all species observed post-FMT (i.e. “fraction of post-FMT strain populations attributable to recipient baseline strains”).

*Coexistence index*: average fraction of coexisting donor and recipient strains (defined as in table S5) among all species post-FMT.

*Colonisation index*: average fraction of donor strains among all species post-FMT.

*Resilience index*: average fraction of persistent recipient strains among species present in the recipient at pre-FMT baseline (i.e. “fraction of recipient baseline strain population that persists post-FMT”).

*Invasiveness index*: average fraction of colonising donor strains among species present in the donor at baseline (i.e. “fraction of donor strain population that successfully colonises”).

As defined above, the persistence, coexistence and colonisation indices describe how much of the post-FMT strain population is attributable to the recipient, the donor, or to coexistence. In contrast, the resilience and invasiveness indices describe how much of the baseline recipient and donor strain populations are carried over post FMT.

### Modeling and predicting strain-level FMT outcomes

We explored a large set of covariates as putative predictor variables for FMT outcomes, grouped into the following categories: (i) host clinical and procedural variables (e.g. FMT indication, pre-FMT bowel preparation, FMT route, etc.); (ii) community-level taxonomic diversity (species richness, community composition, etc.); (iii) community-level metabolic profiles (abundance of specific pathways); (iv) abundance profiles of individual species; (v) strain-level outcomes for other species in the system; and (vi) focal species characteristics, including strain-level diversity; see table S6 for a full list of covariates and their definitions. We further classified covariates as either predictive *ex ante* variables (i.e. knowable before the FMT is conducted) or *post hoc* variables (i.e. pertaining to the post-FMT state, or the relation between pre- and post-FMT states).

We built two types of models to predict FMT strain-level outcomes based on these covariates: (i) FMT-wide models, using summary outcome metrics across all species in a time series (persistence index, colonisation index, etc.; see above) as response variables; and (ii) per-species models for 159 species observed in ≥50 FMTs, using each species’ strain-level outcome in every scored time series as response variable. Unless otherwise indicated, the last available time point for each FMT time series was used. Models were built for each covariate category separately, as well as for combinations of all *ex ante* and all *post hoc* variables, respectively.

Given that the number of covariates greatly exceeded the number of available FMT time series, and that several covariates were correlated with each other (see Fig S3), FMT outcomes were modelled using ten times 5-fold cross-validated LASSO-regularised regression, as implemented in the R package *glmnet* ^88^. Regression coefficients were chosen at one standard error from the cross-validated minimum lambda value and averaged across validation folds.

Linear LASSO regression was used to model outcomes with continuous response variables, both for FMT-wide outcomes (persistence index, etc.) and for the fraction of colonising, persisting and coexisting strains per species across FMTs. For linear models, R^2^ of predictions on test sets was averaged across validation folds. Moreover, logistic LASSO regression was used to additionally model binarised FMT outcomes per species, defined as recipient strain resilience, recipient strain turnover, and donor strain takeover, based on further summarising outcome categories in table S5. For logistic models, accuracy was assessed as Area Under the Receiver Operating Characteristic curve (AUROC), averaged across validation folds.

### Data availability

Raw metagenomic sequencing data have been uploaded to the European Nucleotide Archive under the accession numbers PRJEB46777, PRJEB46778, PRJEB46779 and PRJEB46780. Contextual data is available as online supplementary material. Metagenome-assembled genomes are available for download via Zenodo (DOI 10.5281/zenodo.5534163).

## Acknowledgements

We thank Alan Moss (Harvard U) and Casey Morrow (U Alabama Birmingham) for providing additional information on the US_CD and US_rCDI_2 cohorts used in this study. We further thank Renato J Alves, Anna Schwartz, Michael Kuhn, Pamela Ferretti, Sofia K Forslund, and members of the Bork lab at EMBL for support and constructive discussions.

## Figure & Table Legends

**Figure S1**. Strain-level outcomes for 159 prevalent species tracked across FMT time series. FMTs (columns) are organised by indication, scope (allogenic vs autologous) and clinical success. Species (rows) are organised by underlying phylogeny, corresponding to that shown in Fig 3A.

**Figure S2**. Ternary plots of the post-FMT strain population space for each of 159 prevalent species. Each dot corresponds to a *conspecific* FMT outcome for the focal species; non-conspecific outcomes (acquisition or loss of entire strain population, complete failure to colonise) are not shown.

**Figure S3**. Collinearity of relevant predictor variables. The heatmap shows pairwise Spearman correlation values between candidate predictor variables, organised by type, scope & resolution as outlined in table S6.

**Table S1**. Overview of datasets used in this study.

**Table S2**. Contextual information for all 142 analysed FMT time series.

**Table S3**. Contextual information for all 556 analysed faecal metagenome samples.

**Table S4**. Contextual data for 1,089 species included in this study.

**Table S5**. Strain-level FMT outcome scoring rules, based on determinant microbial SNVs and gene content per species pan-genome in sample triplets of donor baseline, recipient baseline and recipient post-FMT. See Methods and Fig 1C for details and visualisation.

**Table S6**. Overview and definitions of variables tested for association with FMT outcomes. Variables are categorised by type, scope (pertaining to the host or microbiome, to the donor or recipient) and resolution (host-level, community-level, species-level, strain-level). See Methods for additional details.

**Table S7**. mOTU taxonomic profiles.

**Table S8**. GMM profiles.

## Notes

### Competing Interest Statement

TJB has a pecuniary interest in the Centre for Digestive Diseases, is a medical advisor to Finch therapeutics and holds patents in FMT treatment. CP received grant support from Takeda, Gilead, and Perspectum, consultancy fees from Shire and Pliant, and speaker's fees from Tillotts and Takeda. MN and WdV are founders and members of the Scientific Advisory Board of Caelus Health, the Netherlands. MN is a Scientific Advisory Board member of Kaleido Biosciences, USA. WdV is founder and Scientific Advisory Board member of A-mansia Biotech Belgium.
None of these conflicts bear any relevance to the content of the current manuscript.

