## Supplementary figures and images for "Drivers and Determinants of Strain Dynamics Following Faecal Microbiota Transplantation"

### Figure S1

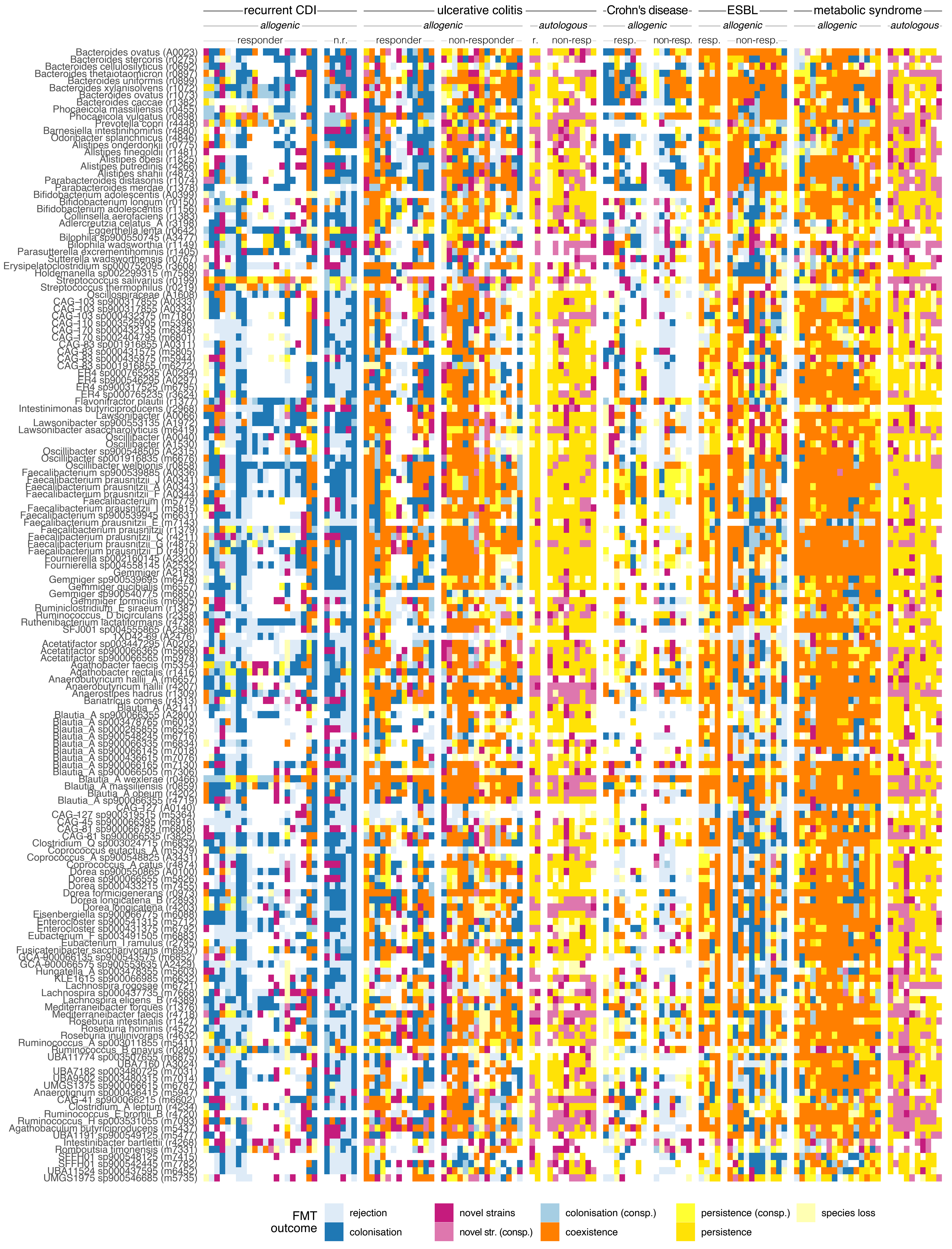

### Figure S2

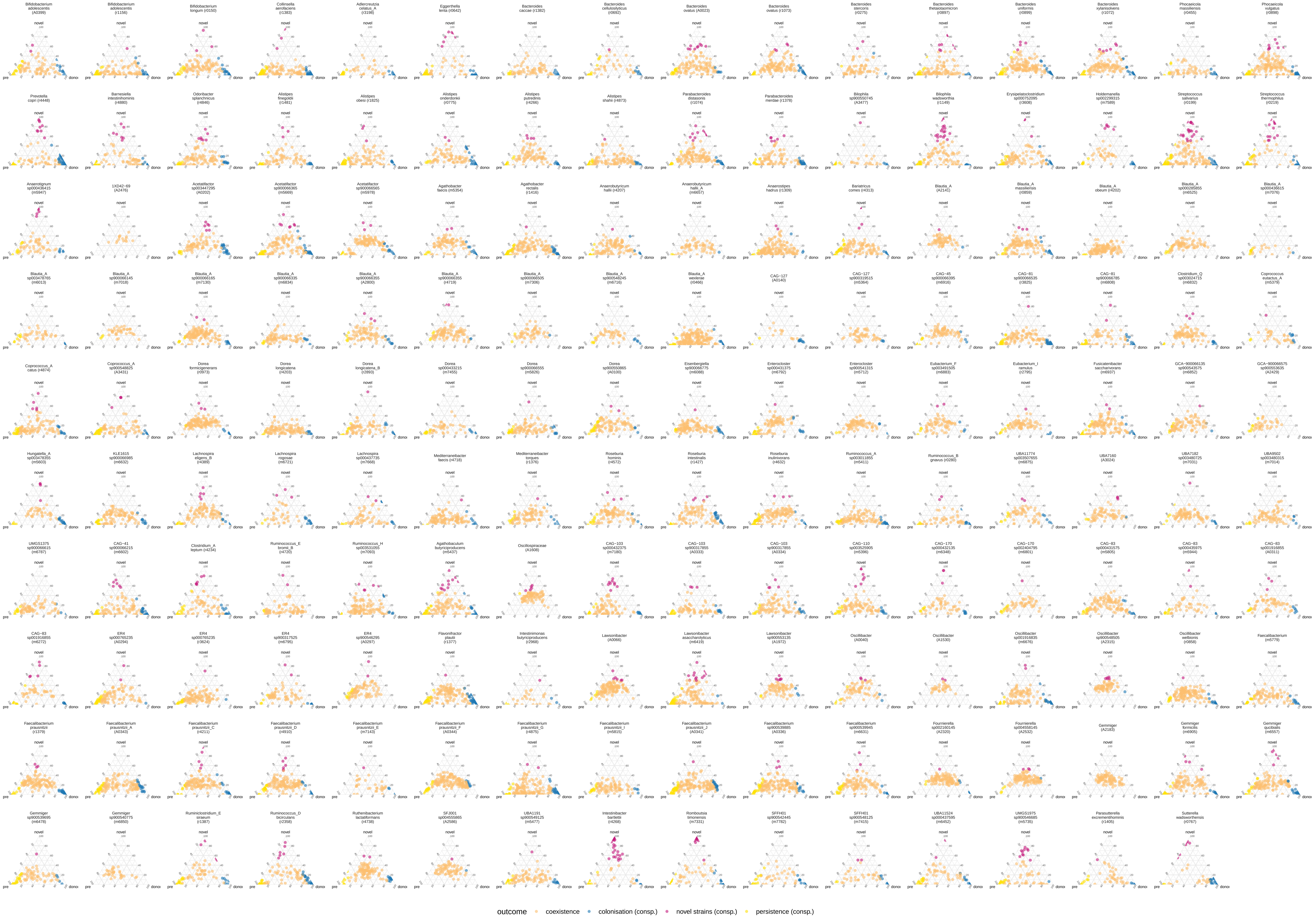

### Figure S3

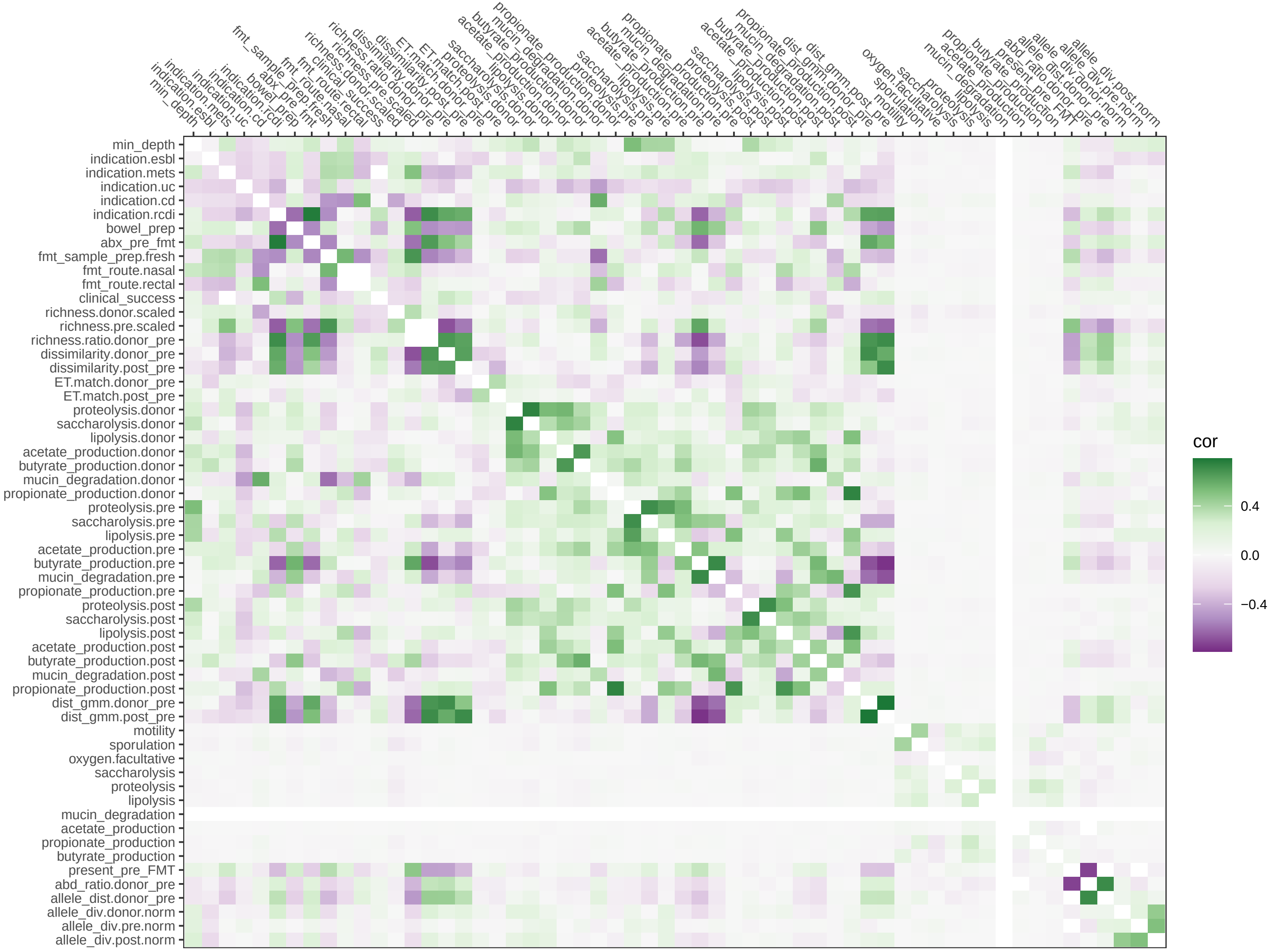
